# IL-27 receptor signaling regulated stress myelopoiesis drives Abdominal Aortic Aneurysm development

**DOI:** 10.1101/555581

**Authors:** luliia O. Peshkova, Turan Aghayev, Aliia R. Fatkhullina, Petr Makhov, Satoru Eguchi, Yin Fei Tan, Andrew V. Kossenkov, Stephen Sykes, Ekaterina K. Koltsova

**Affiliations:** Blood Cell Development and Function Program, Fox Chase Cancer Center, Philadelphia, PA, 19111, USA; Cancer Biology Program, Fox Chase Cancer Center, Philadelphia, PA, 19111, USA; Lewis Katz School of Medicine, Temple University, Cardiovascular Research Center, Philadelphia, PA, 19140, USA; Genomics Facility, Fox Chase Cancer Center, Philadelphia, PA, 19111, USA; Bioinformatics Facility, The Wistar Institute, Philadelphia, PA 19104, USA

**Author notes:** These authors equally contributed to this work (luliia O. Peshkova, Turan Aghayev). Address correspondence to: Ekaterina Koltsova, MD, PhD, Blood Cell Development and Function Program, Fox Chase Cancer Center, P2151.

**Keywords:** cytokines, myeloid cells, hematopoietic stem and progenitor cells, abdominal aortic aneurysm

## Abstract

Abdominal Aortic Aneurysm (AAA) is a vascular disease, where aortic wall degradation is mediated by accumulated immune cells. Though cytokines regulate the inflammatory milieu within the aortic wall, their contribution to AAA through distant alterations, particularly in the control of hematopoietic stem cells proliferation and myeloid cell differentiation remains poorly defined. Here we report an unexpected pathogenic role for interleukin-27 receptor (IL-27R) in AAA development as genetic inactivation of IL-27R protected mice from AAA induced by Angiotensin (Ang) II. The mitigation of AAA in IL-27R deficient mice is associated with a blunted accumulation of myeloid cells in suprarenal aortas due to the surprising attenuation of Ang II-induced expansion of HSCs. The loss of IL-27R engages transcriptional programs that promote HSCs quiescence and suppresses myeloid lineage differentiation, decreasing mature cell production and myeloid cell accumulation in the aorta.

We, therefore, illuminate how a prominent vascular disease can be distantly driven by cytokine dependent regulation of the bone marrow precursors.

## Introduction

Abdominal Aortic Aneurysm (AAA) is a cardiovascular disease, which due to limited therapeutic options is a significant cause of death in the elderly population^1^. AAA is characterized by immune cell infiltration into the aortic wall and progressive degradation of the medial layer, resulting in the dilatation and rupture of the aorta, ultimately leading to fatal bleeding^2,3,4^. Since current standard of care for AAA is still limited to surgical interventions at the late stages, a better understanding of its mechanisms, particularly the inflammatory nature of this disease, is urgently needed. Various risk factors are associated with AAA pathogenesis, including elevated blood pressure that is mediated by the activation of the renin-angiotensin system (RAS) and an increase in Angiotensin II (Ang II). Long-term infusion of Ang II in mice is able to recapitulate many aspects of AAA in humans, as it induces AAA formation in the ascending aorta^5,6^. In addition to regulating blood pressure, Ang II is known to control the function of various vascular wall cells predominantly through AT1a receptor (AT1aR) signaling^7,8,9,10,11,12^.

Chronic inflammation caused by the infiltration of various immune cells is a key driver of AAA etiology and pathogenesis^2,3,4^. In particular, neutrophils and monocytes, which are bone marrow (BM) derived myeloid cells, play an important role in the initiation of aortic wall destruction^13,14,15,16,17^. Their output by hematopoietic stem and progenitors cells (HSPCs) in the BM can be influenced by environmental stimuli and stresses. During infections or inflammatory insult, a variety of cytokines and danger associated molecular patterns (DAMPs) stimulate HSPCs to rapidly increase the production of innate immune cells^18,19,20^. Although alterations in myelopoiesis have been reported to play a role in AAA progression^4,16^, the key cytokine signaling factors that control HSPCs fate and myelopoiesis in AAA have yet to be determined. Hematopoietic stem cells (HSCs) were shown to express the Angiotensin II receptor, AT1aR and respond to elevated Ang II levels during cancer development^21^. Furthermore, Ang II infusion promotes the expansion of murine HSPCs, suggesting that HSPCs may be affected during AAA pathogenesis characterized by elevated levels of Ang II^22^.

Interleukin (IL)-27 is a member of IL-6/IL-12 cytokine superfamily that regulates various hematopoietic and non-hematopoietic cells in infectious diseases and autoimmunity^23,24,25,26,27,28^. Nevertheless, knowledge regarding the role of IL-27 in regulation of hematopoiesis in chronic inflammatory diseases remains limited. Moreover, the role of IL-27R signaling in AAA pathogenesis, inflammation and HSPCs homeostasis has yet to be investigated.

Here we found that inactivation of IL-27R protects mice from Ang II-induced AAA. Mechanistically, we showed that IL-27R signaling is essential to drive Ang II-mediated HSPCs proliferation and myeloid differentiation. Our unexpected findings illustrate how IL-27 signaling acts distantly to control AAA development by cooperating with stress-induced factors in the bone marrow to accelerate stress myelopoiesis, promote the production of myeloid cells that are subsequently recruited to the aortic wall mediating its destruction.

## Results

### IL-27R signaling promotes Ang II-induced abdominal aortic aneurysm development

Inactivation of IL-27R exacerbates atherosclerosis^26,27,28^ and leads to the development of abdominal aortic lesions which typically are rare in atherosclerotic mice. As atherosclerosis and AAA may share some underlying chronic inflammatory mechanisms^29^, we anticipated that IL-27R-deficiency would increase inflammation and promote AAA.

To evaluate the potential role of IL-27R signaling in AAA, we employed a well characterized mouse model of AAA driven by chronic Ang II infusion through a surgically implanted osmotic mini-pump (800 ng · kg^−1^ min^−1^) into mice on *Apoe*^−/−^ background^5,30^. To exclude any differences in genetics or microbiota, we used cage-mate and littermate controls. Male and female *Apoe*^−/−^, *Apoe*^−/−^*Il27ra*^+/−^ or *Apoe*^−/−^*Il27ra*^−/−^ mice were fed with a western diet (WD) for 8 weeks followed by Ang II pump implantation. Four weeks later, mice were assessed for bulging of abdominal aorta and AAA development (Figure 1A). We found that Ang II infusion induces AAA formation in IL-27R proficient *Apoe*^−/−^ and *Apoe*^−/−^*Il27ra*^+/−^ control mice, while unexpectedly the incidence of AAA was markedly reduced in *Apoe*^−/−^*Il27ra*^−/−^ mice (Figure 1B-F). Both male and female *Apoe*^−/−^ and *Apoe*^−/−^*Il27ra*^+/−^ mice developed larger AAAs with visual hemorrhages in the artery walls compared to their *Apoe*^−/−^*Il27ra*^−/−^ counterparts (Figure 1B, C). Blood pressure was elevated in response to Ang II infusion; but IL-27R regulated AAA independent of effects on blood pressure, body weight or plasma lipids (Supplemental Figure 1A,B, not shown). Verhoff-van Gieson staining revealed extensive disruption and degradation of elastic lamina in the aortas of both *Apoe*^−/−^ and *Apoe*^−/−^*Il27ra*^+/−^ mice, while no significant elastin degradation was found in *Apoe*^−/−^*Il27ra*^−/−^ mice (Figure 1D). Female *Apoe*^−/−^ and *Apoe*^−/−^*Il27ra*^+/−^ mice (Figure 1E) developed slightly lower rates of AAA than did their male counterparts (Figure 1F); however, the incidence of AAA was reduced by IL-27R-deficiency in *Apoe*^−/−^*Il27ra*^−/−^ mice of both genders (Figure 1E,F), indicating that IL-27R mediated promotion of AAA in both genders. While both *Apoe*^−/−^ and *Apoe*^−/−^*Il27ra*^+/−^ mice experienced significant AAA-related mortality, *Apoe*^−/−^*Il27ra*^−/−^ mice exhibited a 100% (Figure 1G, H). Pathological severity index based on the level of aortic wall degradation and immune infiltrate as previously described^31^, revealed that both female and male *Apoe*^−/−^ and *Apoe*^−/−^*Il27ra*^+/−^ mice had more advanced AAA (IV stage), than was observed in IL-27R deficient *Apoe*^−/−^ mice, where AAA progression was restricted to the early stages (I-II) (Supplemental Figure 2). Therefore, our data demonstrate that IL-27R signaling unexpectedly promotes AAA.

**Figure 1.**
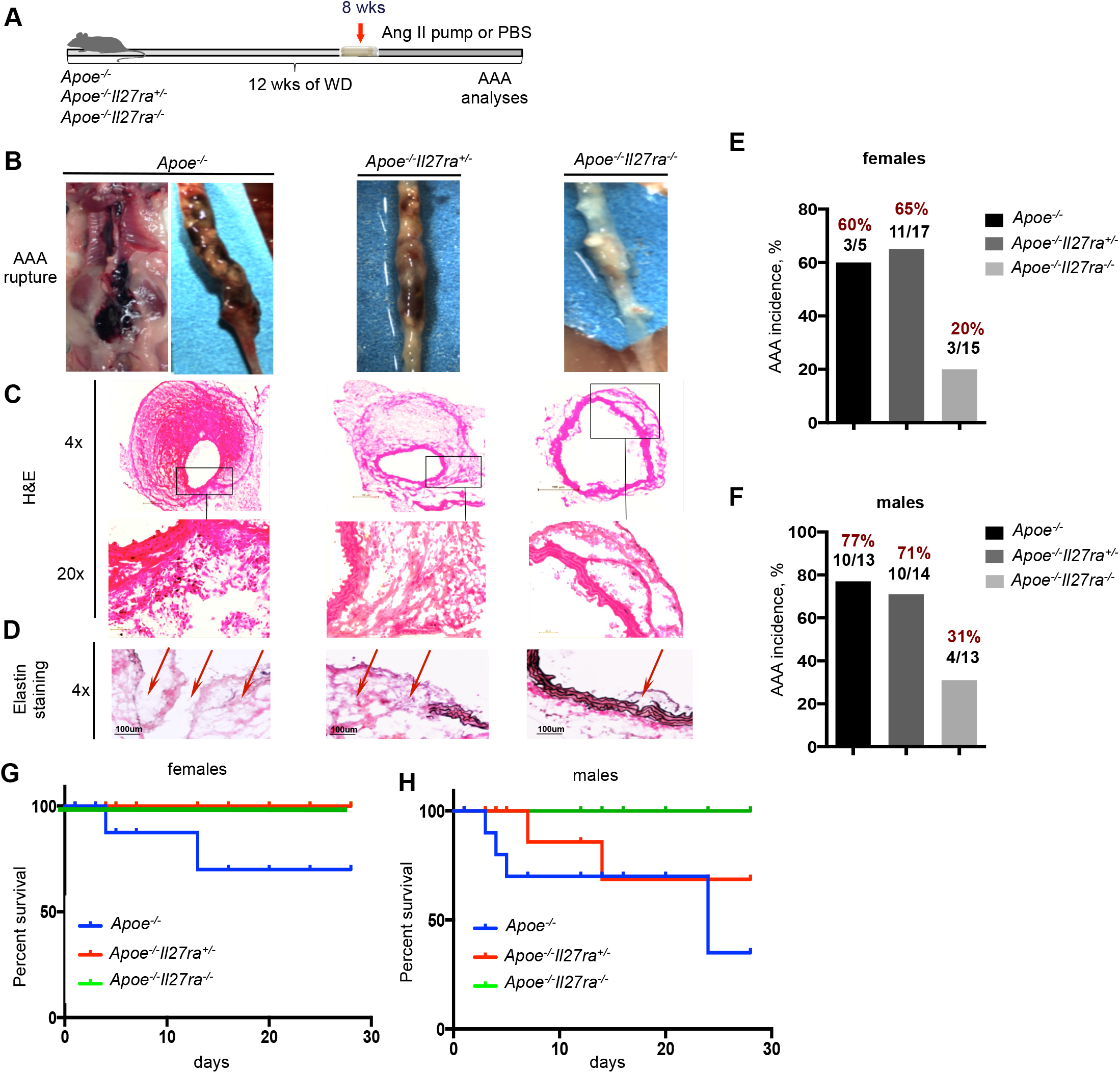
IL-27R deficiency protects from Ang-II-induced AAA development. **(A)** Scheme of the experiment. *Apoe*^−/−^, *Apoe*^−/−^*Il27ra*^+/−^ and *Apoe*^−/−^*Il27ra*^−/−^ female and male mice were fed with WD for over-all period of 12 weeks, where last 4 weeks of feeding they were implanted with pumps containing Ang II or PBS. **(B)** Representative images of suprarenal aortas with developed AAA. **(C)** Hematoxylin and eosin (H&E) staining, **(D)** Verhoeff-Van Gieson (EVG) staining of frozen sections of AAA from *Apoe*^−/−^, *Apoe*^−/−^*Il27ra*^+/−^ or *Apoe*^−/−^*Il27ra*^−/−^ mice after Ang II infusion. Scale bars, 100 μm. Black-elastin, red-collagen, blue-nuclei. Arrows indicate ruptured elastic lamina. Percentage of AAA incidence among *Apoe*^−/−^ (n=5), *Apoe*^−/−^*Il27ra*^+/−^ (n=17) and *Apoe*^−/−^*Il27ra*^−/−^ (n=15) female **(E)** and *Apoe*^−/−^ (n=13), *Apoe*^−/−^*Il27ra*^+/−^ (n=14) and *Apoe*^−/−^*Il27ra*^−/−^ (n=13) male mice **(F)**. Survival curves for *Apoe*^−/−^ (2 out of 5 died), *Apoe*^−/−^*Il27ra*^+/−^ (0 out of 17 died) and *Apoe*^−/−^*Il27ra*^−/−^ (0 out of 15 died) female **(G)** and *Apoe*^−/−^ (4 out of 13 died), *Apoe*^−/−^ *Il27ra*^+/−^ (2 out of 14 died) and *Apoe*^−/−^*Il27ra*^−/−^ (0 out of 13 died) male mice **(H)** during 28 days of Ang II infusion.

### *Apoe*^−/−^*Il27ra*^−/−^ mice exhibit reduced accumulation of myeloid cells in suprarenal aortas

AAA progression is associated with increased accumulation of various immune cells at the site of vessel injury^2,3,4^. To characterize immune cell accumulation, isolated suprarenal aortas (with or without AAA) were digested and analyzed by FACS. FACS analysis revealed a significant reduction in the percentage and number of hematopoietic CD45^+^ cells in suprarenal aortas of *Apoe*^−/−^*Il27ra*^−/−^ mice compared to *Apoe*^−/−^*Il27ra*^+/−^ controls (Figure 2A). Among CD45^+^ cells, the number of CD11b^+^, CD11b^+^CD11c^+^ and CD11c^+^ myeloid cell subsets were also significantly diminished in *Apoe*^−/−^*Il27ra*^−/−^ (Figure 2B). We observed a striking reduction in different monocyte subsets (Ly6C^high^ and Ly6C^low^) as well as neutrophils (Ly6G^+^) in AAA lesions of *Apoe*^−/−^*Il27ra*^−/−^ mice compared to *Apoe*^−/−^*Il27ra*^+/−^ mice (Figure 2C). Immunofluorescence staining of isolated AAAs further confirmed limited accumulation of CD11b^+^ myeloid cells and particularly Ly6G^+^ neutrophils in adventitia of AAA of *Apoe*^−/−^*Il27ra*^−/−^ mice (Figure 2D). Q-RT-PCR analysis revealed the reduction in expression of molecules involved in attraction and tissue trafficking of myeloid cells, such as chemokines (CCL2, CCL5, CCL22) and cytokines (TNF-α, IL-1β) as well as matrix metalloproteinases (MMPs) (Figure 2E-G).

**Figure 2.**
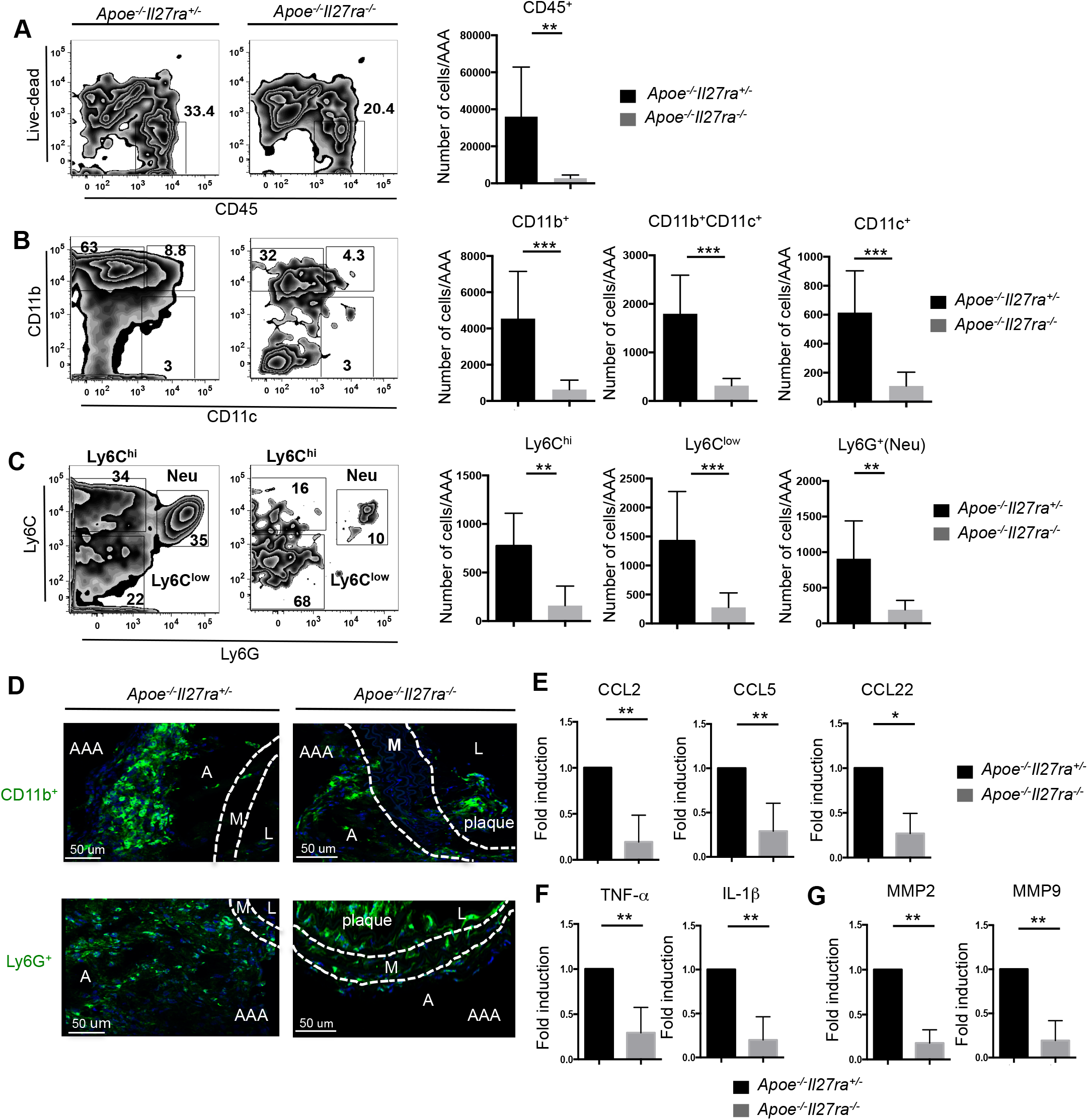
Limited accumulation of immune cells in AAA of IL-27R deficient mice. Singlecell suspensions were obtained from suprarenal aortas/AAA of *Apoe*^−/−^*Il27ra*^+/−^ or *Apoe*^−/−^*Il27ra*^−/−^ (n=6) mice fed with WD and infused with Ang II. Cells were stained for live/dead, CD45, CD11b, CD11c, Ly6G and Ly6C and analyzed by FACS. **(A, left)** Representative dot plots of CD45^+^ cells. Numbers indicate percentage in each quadrant, **(A, right)** Absolute number of live CD45^+^ cells. **(B)** Representative dot plots and absolute number of CD45^+^ live CD11b^+^, CD11b^+^CD11c^+^ and CD11c^+^ cells. **(C)** Representative dot plots and absolute number of live CD11b^+^Ly6C^hi^, CD11b^+^Ly6G^+^ (Neu) and CD11b^+^Ly6C^low^ cells. **(D)** Localization and abundance of CD11b^+^ and Ly6G^+^ (Neu) cells in suprarenal aortas/AAA of *Apoe*^−/−^*Il27ra*^+/−^ and *Apoe*^−/−^*Il27ra*^−/−^ mice infused with Ang II as seen by immunofluorescence using Leica SP8 confocal microscope. L-lumen, M-media, A-adventitia, AAA-abdominal aortic aneurysm. Representative images from 3 independent experiments. Relative gene expression of chemokines **(E)**, cytokines **(F)** and MMPs **(G)** and in suprarenal aortas/AAA of *Apoe*^−/−^*Il27ra*^−/−^ (n=6) mice were normalized to L-32 gene expression and then normalized to gene expression in suprarenal aortas/AAA of control *Apoe*^−/−^*Il27ra*^+/−^ (n=8) mice. *p<0.05. **p<0.01, ***p<0.005. Data are mean ± SEM from at least 3 independent experiments.

Thus, IL-27R deficiency leads to the reduced accumulation of myeloid cells, especially monocytes and neutrophils in suprarenal aortas. This further results in a decreased expression of myeloid-derived chemokines, cytokines and enzymes in the area of AAA formation.

### IL-27R signaling is required for the regulation of BM HSPCs during AAA development

Inflammation associated with infection and injury is also known to modulate output and mobilization of myeloid cells from the bone marrow^20,32^. Next we sought to determine if reduced myeloid cell accumulation in AAA sites of suprarenal aortas and AAA development in the absence of IL-27R is associated with alterations in hematopoiesis. We analyzed the cellular composition of BM isolated from *Apoe*^−/−^*Il27ra*^−/−^and control *Apoe*^−/−^*Il27ra*^+/−^ mice. In steady-state (PBS-infusion), IL-27R-deficiency did not cause dramatic changes in LSK compartment (Lineage^−^, Sca-1^+^, c-Kit^+^) or in myeloid progenitors (Lineage^−^, c-Kit^+^, Sca-1^−^) (Figure 3A, C, D), including common myeloid progenitors (CMP, Lineage^−^, Sca-1^−^, c-Kit^+^, FcgII/III^−^, CD34^+^) and granulocyte-monocyte progenitors (GMP, Lineage^−^, Sca-1^−^, c-Kit^+^, FcgII/III^+^, CD34^+^) (Figure 3D and data not shown). The percentage of long-term progenitors (LT-HSC, Lineage^−^, Sca-1^+^, c-Kit^+^, CD150^+^, CD48^−^) was slightly elevated, while the percentage of LSK CD48^+^CD150^−^ population was slightly reduced in IL-27R deficient mice (Figure 3B,C). In agreement with previous observations^22^, Ang II infusion significantly increased the percentage of LSK and total myeloid progenitors in *Apoe*^−/−^*Il27ra*^+/−^ mice (Figure 3A, C, D) compared to PBS treated “non-stressed” controls, indicating that Ang II-induced AAA development is indeed accompanied by significant changes in myeloid cell development. Surprisingly, Ang II driven expansion of LSK or myeloid precursors was blunted in the absence of IL-27R (Figure 3A, C, D). The percentage of LT-HSC did not change in response to Ang II in *Apoe*^−/−^*Il27ra*^+/−^ but remained elevated in *Apoe*^−/−^*Il27ra*^−/−^ mice (Fig 3B, C). A mild increase in CD48^+^CD150^−^ population in response to Ang II was found in controls, while Ang II-treated IL-27R deficient mice still displayed a reduction of these cells (Figure 3B, C). Ang II infusion also decreased CMP and GMP in *Apoe*^−/−^*Il27ra*^−/−^ (Figure 3D). Only a slight reduction of myeloid progenitors in the absence of IL-27R signaling was found in the spleen (data not shown).

**Figure 3.**
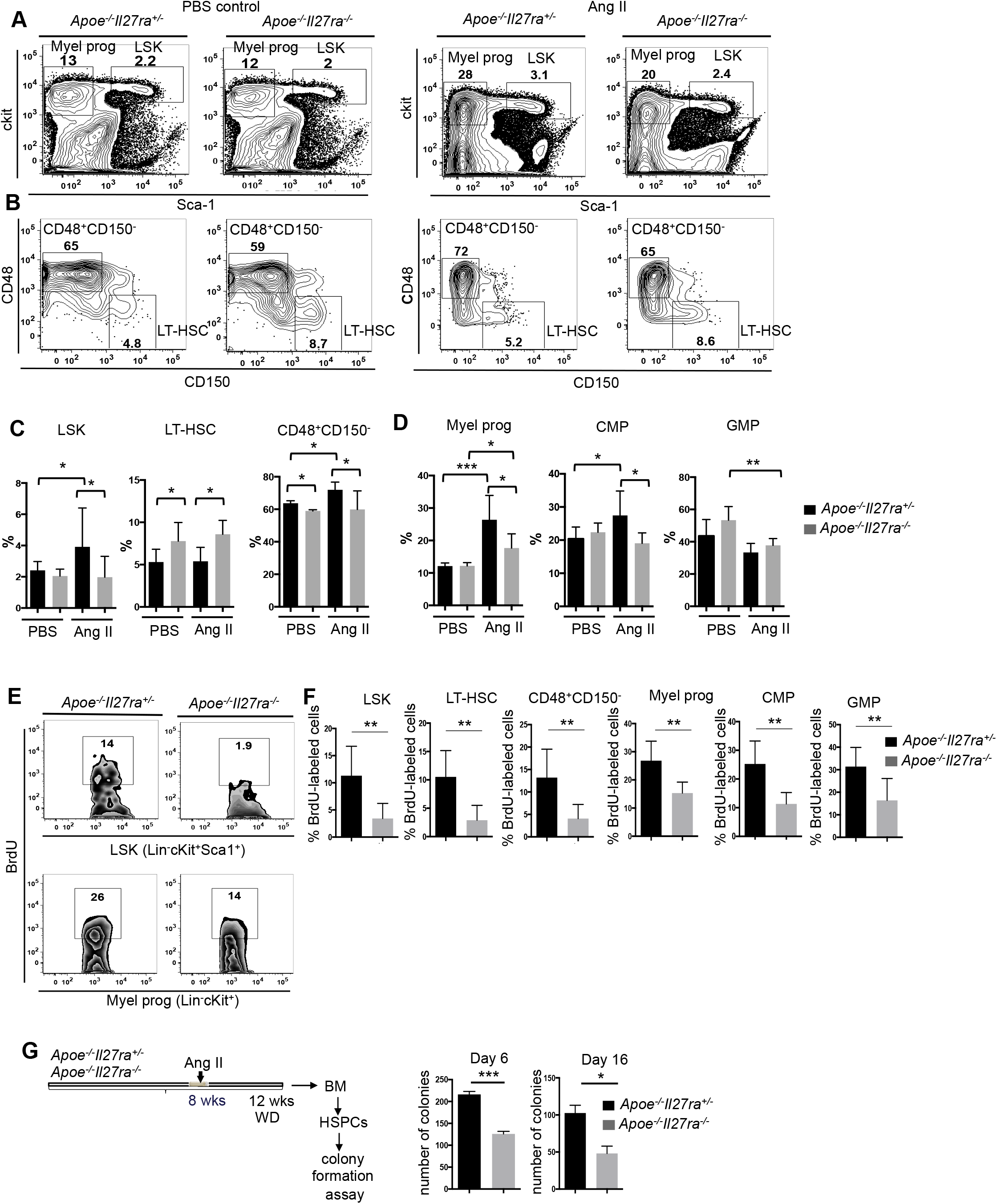
Limited expansion of HSCs and myeloid progenitors in bone marrow of *Apoe*^−/−^ *Il27ra*^−/−^ mice in response to Ang II infusion. Bone marrow was obtained from *Apoe*^−/−^*Il27ra*^+/−^ (n=7-11) and *Apoe*^−/−^*Il27ra*^−/−^ (n=6-12) mice fed with WD for over-all period of 12 weeks where last 4 weeks of feeding they were implanted with pumps containing PBS or Ang II. **(A)** Representative dot plots of live LSK (Lin^−^c-kit^+^Sca1^+^) and myeloid progenitors (Lin^−^c-kit^+^Sca1^−^). **(B)** Representative dot plots of live LT-HSC (CD150^+^CD48^−^) and CD48^+^CD150^−^ cells. Average percentage of LSK, LT-HSC and CD48^+^CD150^−^ cells **(C)**, myeloid progenitors, CMP (Lin^−^c-kit^+^CD34^+^FcgR^−^) and GMP (Lin^−^c-kit^+^CD34^+^FcgR^+^) **(D)**. Data are mean ± SEM from 3 independent experiments. Proliferation of bone marrow HSPCs isolated from *Apoe*^−/−^*Il27ra*^+/−^ (n=8) or *Apoe*^−/−^*Il27ra*^−/−^ (n=7) mice fed with WD and infused with Ang II as determined by BrdU incorporation. Representative dot plots **(E)** and percentage **(B)** of live BrdU positive LSK (Lin^−^c-kit^+^Sca1^+^) and myeloid progenitors (Lin^−^c-kit^+^Sca1^−^), including LT-HSC (CD150^+^CD48^−^), CD48^+^CD150^−^, CMP (Lin^−^c-kit^+^CD34^+^FcgR^−^) and GMP (Lin^−^c-kit^+^CD34^+^FcgR^+^). Data are mean ± SEM from 2 independent experiments. (G) Scheme of *ex vivo* colony formation experiment. Lin^−^ HSPCs were isolated from bone marrow of *Apoe*^−/−^*Il27ra*^+/−^ (n=5) or *Apoe*^−/−^*Il27ra*^−/−^ (n=5) mice infused with Ang II and plated in M3434 media under myeloid conditions. Colony formation was accessed on days 6 and 16. Representative data are mean ± SEM from 2 independent experiments. *p<0.05, **p<0.01, ***p<0.005

The Ang II receptor, AT1aR is expressed on HSPCs^21^ and Ang II infusion promotes both proliferation and myeloid-biased differentiation of HSCs^22^. To test the ability of HSPCs to proliferate in response to Ang II, we assessed BM cell proliferation by *in vivo* BrDU incorporation. We found that BrDU incorporation was significantly lower among various precursor cells in the BM of Ang II-infused *Apoe*^−/−^*Il27ra*^−/−^ mice compared to *Apoe*^−/−^*Il27ra*^+/−^ controls (Figure 3E, F), while no difference in proliferation level was observed in PBS-infused mice (Supplemental Figure 3). These data suggest that IL-27R signaling potentiates the proliferative effect of Ang II on HSPCs.

To evaluate the effect of IL-27R signaling on the clonogenic and differentiation capacities of HSPCs, we performed *ex vivo* colony formation assay. Lineage-depleted (Lin-) BM HSPCs were isolated from *Apoe*^−/−^*Il27ra*^−/−^ and *Apoe*^−/−^*Il27ra*^+/−^ mice after 4 weeks of Ang II infusion and cultured in M3434 media under myeloid differentiation conditions. We found that IL-27R deficient HSPCs showed a marked decrease in total numbers of colonies compared to IL-27R sufficient cells (Figure 3G). Our data suggest that IL-27R signaling potentiates the Ang II-induced “stress” myelopoiesis during AAA development.

### IL-27R deficient HSCs are less competitive in stress-induced expansion and fail to induce AAA

To evaluate if a cell-intrinsic lack of IL-27R signals render HSPCs unable to expand and contribute to AAA in an overall IL-27R sufficient environment, we conducted competitive bone marrow transfer (BMT) analysis. BM isolated from *Apoe*^−/−^ CD45.1^+^ (further referred as WT) or *Apoe*^−/−^*Il27ra*^−/−^ CD45.2^+^ (further referred as *Il27ra*^−/−^) congenic mice were mixed in different ratios including 90%:10%, 50%:50% and 10%:90%, respectively, and transplanted into lethally irradiated *Apoe*^−/−^ CD45.1 IL-27R sufficient recipients. Mice were allowed to reconstitute the BM for 4 weeks followed by 10 week WD feeding. The efficiency of BM reconstitution was confirmed by FACS for CD45.1/CD45.2 ratio in the blood of naive unchallenged mice 4 weeks after BMT (Supplemental Figure 4). After 8 weeks of WD feeding mice were implanted with Ang II pump to induce AAA and assess the effect of Ang II on the BM cells and their role in AAA (Figure 4A). Interestingly, all mice transplanted with a 90%wt:10%*Il27ra*^−/−^ mix of BM developed late stages of AAA and some died due to AAA rupture (Figure 4B, C); however, none of the mice transplanted with a 10%wt:90%*Il27ra*^−/−^ mix of BM developed AAA (Figure 4B, C) and 5 mice out of 13 transplanted with mix of 50%wt:50%*Il27ra*^−/−^ BM developed AAA (Figure 4B, C).

**Figure 4.**
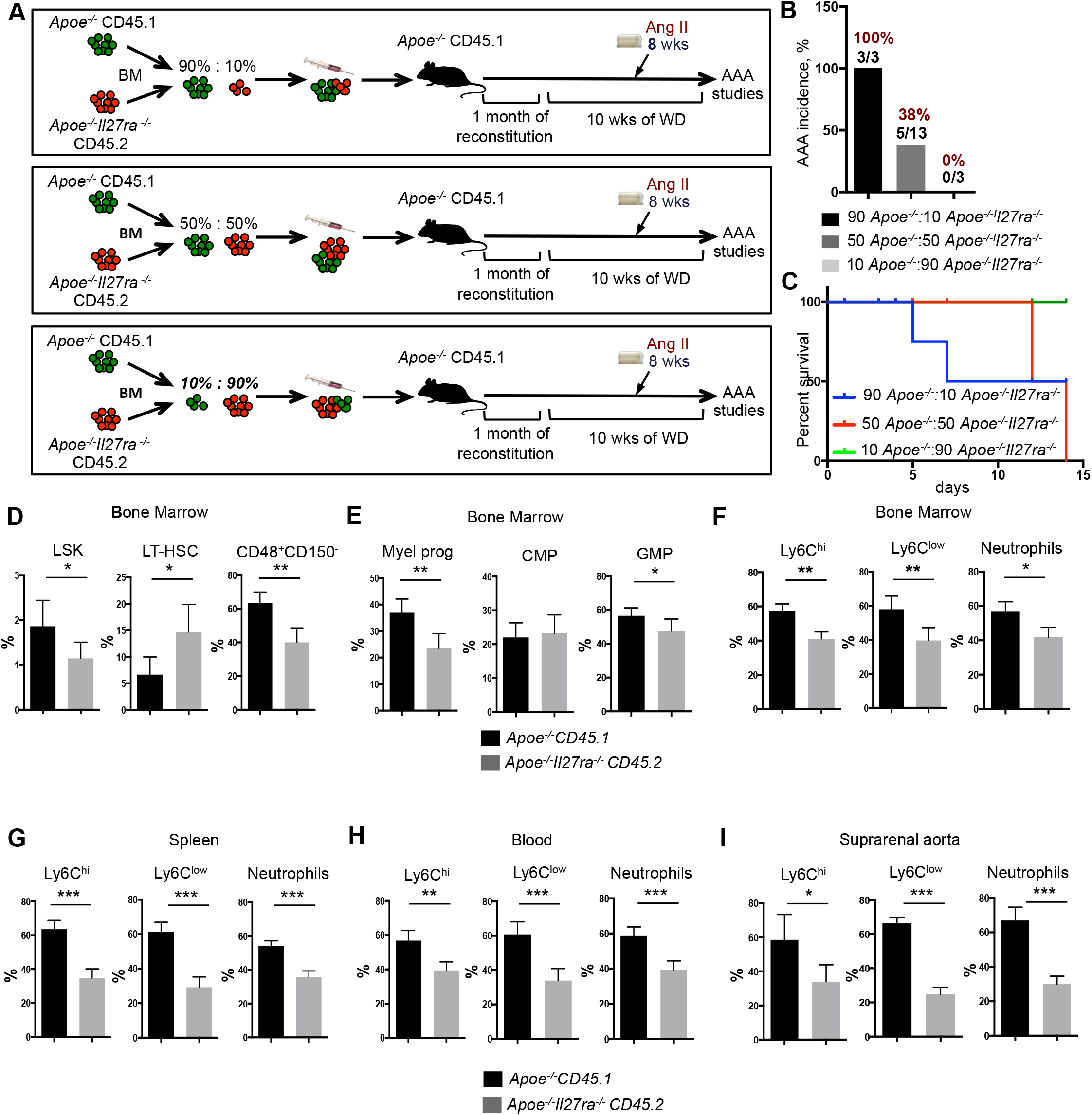
IL-27R deficient hematopoietic stem cells are less competitive in stress induced expansion and defective in induction of AAA. **(A)** Scheme of experiment. *Apoe*^−/−^ CD45.1 recipient mice were lethally irradiated and reconstituted with donor mixes of *Apoe*^−/−^ CD45.1 and *Apoe*^−/−^*Il27ra*^−/−^ CD45.2 total bone marrow cells in a 90%:10%, 50%:50% and 10%:90% ratio. 4 weeks after reconstitution mice were placed on WD for 10 weeks, where during last 2 weeks of feeding they were infused with Ang II. **(B)** Percentage of AAA incidence among *Apoe*^−/−^ CD45.1 recipient mice receiving bone marrow mixture in a ratio: 90% *Apoe*^−/−^:10% *Apoe*^−/−^*Il27ra*^−/−^ (n=3), 50% *Apoe*^−/−^:50% *Apoe*^−/−^*Il27ra*^−/−^ (n=13) or 10% *Apoe*^−/−^:90% *Apoe*^−/−^*Il27ra*^−/−^ (n=3). **(C)** Survival curves for *Apoe*^−/−^ CD45.1 recipient mice receiving bone marrow mixture in a ratio: 90% *Apoe*^−/−^:10% *Apoe*^−/−^*Il27ra*^−/−^ (2 out of 3 died), 50% *Apoe*^−/−^:50% *Apoe*^−/−^*Il27ra*^−/−^ (5 out of 13 died) or 10% *Apoe*^−/−^:90% *Apoe*^−/−^*Il27ra*^−/−^ (0 out of 3 died) during 28 days of Ang II infusion. Proportion of live donor-specific cells in *Apoe*^−/−^ CD45.1 recipient mice receiving bone marrow mix in a ratio 50% *Apoe*^−/−^:50% *Apoe*^−/−^*Il27ra*^−/−^ **(D)** LSK cells, including LT-HSC and CD48^+^CD150^−^, **(E)** myeloid progenitors, including CMP and GMP in bone marrow. Proportion of donor-specific mature Ly6C^hi+^, Ly6C^low+^ monocytes and Ly6G^+^ neutrophils in bone marrow **(F)**, spleen **(G)**, blood **(H)** and suprarenal aorta **(I)**. *p<0.05, **p<0.01, ***p<0.005. Data are mean ± SEM from 2 independent experiments.

Chimeric recipient mice received 50%wt:50% *Il27ra*^−/−^ BM mix and infused with Ang II were used to compare side by side the development and accumulation of IL-27R sufficient and IL-27R deficient myeloid cells in IL-27R sufficient environment. Both WT and *Il27ra*^−/−^ donors cells exhibited ~ 50% reconstitution of peripheral blood 4 weeks after transplantation (Supplemental Figure 4). However, the analysis of HSPCs compartment in the BM of chimeric mice infused with Ang II revealed the reduction of *Il27ra*^−/−^ LSK, CD48^+^CD150^−^ and more mature myeloid progenitors compared to WT derived cells within the same mouse (Figure 4D, E). Conversely, the percentage of LT-HSCs derived from *Il27ra*^−/−^ BM was elevated (Figure 4D). In line with limited competitive expansion of IL-27R deficient precursors, the percentage of mature *Il27ra*^−/−^ *Apoe*^−/−^ (CD45.2) derived monocytes and neutrophils were also diminished in the BM, spleen and blood in comparison with IL-27R sufficient *Apoe*^−/−^ (CD45.1) cells from the same recipient (Figure 4F-H). The analysis of suprarenal aortas/AAA revealed a strong preferential accumulation of monocytes and neutrophils derived from WT (IL-27R sufficient) progenitors, while the percentage of accumulated cells derived from *Il27ra*^−/−^ BM was diminished (Figure 4I).

Of note, IL-27R deficiency was previously reported to promote atherosclerosis in *Apoe*^−/−^ mice^28^. Here we found that Ang II infusion into mice with already developed atherosclerosis (8 weeks of WD) strongly accelerated the disease in control, but not in IL-27R deficient mice (Supplemental Figure 5A-C). The analysis of myeloid cells accumulation in 50%wt:50%*Il27ra*^−/−^ BM chimeric mice revealed that monocytes and neutrophils in the aortic arch were mostly cells of wild type *Apoe*^−/−^ (CD45.1) origin, while *Apoe*^−/−^*Il27ra*^−/−^ (CD45.2) cells were significantly underrepresented (Supplemental Figure 5D). These results provide an explanation for equal atherosclerotic lesion sizes between *Apoe*^−/−^*Il27ra*^+/−^ and *Apoe*^−/−^*Il27ra*^−/−^ Ang II infused mice.

Overall, our data suggest that Ang II-driven expansion of HSCs and myelopoiesis facilitates myeloid cell accumulation and AAA and this process is dependent on IL-27R signaling, which provides HSCs and myeloid progenitors with competitive fitness required for the expansion.

### IL-27R signaling regulates HSCs quiescence transcriptional programs in Ang II-induced myelopoiesis

To gain insights into the mechanisms by which IL-27R signaling influences HSCs function and “stress-induced” myelopoiesis in AAA, we performed whole transcriptome RNA sequencing analysis of FACS-sorted CD150^+^CD48^−^ LT-HSCs isolated from the BM of *Apoe*^−/−^*Il27ra*^−/−^, *Apoe*^−/−^*Il27ra*^+/−^ or *Apoe*^−/−^ fed with WD and infused with PBS or Ang II for the last 2 weeks of WD feeding. Consistent with our functional data *in vivo* and *ex vivo*, we found that the lack of IL-27R signaling does not significantly affect transcriptional profile of LT-HSCs in “steady state” (PBS-infused mice), where only 16 genes were differentially expressed (FDR<5%) between *Apoe*^−/−^ *Il27ra*^−/−^ and *Apoe*^−/−^ LT-HSCs; however, Ang II infusion dramatically changed transcriptional profile and led to significant differential expression of 587 genes (FDR<5%) between *Apoe*^−/−^ *Il27ra*^−/−^ and *Apoe*^−/−^ LT-HSCs (Figure 5A). Interestingly, *Agr2*, an inhibitor of p53 pathway^33^ that is downregulated in *Apoe*^−/−^*Il27ra*^−/−^ LT-HSCs, was the only gene that was differentially expressed in both PBS and Ang II-treated IL-27R-deficient HSCs (Figure 5A). Gene set enrichment analysis using Ingenuity Pathway Analysis (IPA) of genes specific to *Apoe*^−/−^*Il27ra*^−/−^ LT-HSCs revealed several significantly de-regulated pathways that had a common effect among all member genes (Supplemental Table 1). Using IPA upstream regulator analysis, we found 24 regulators with a significant number of their known targets (at least 5 targets, p<0.05) overrepresented in the list of significantly affected by IL-27R deficiency genes whose combined behavior showed change in activation status of upstream regulator (Figure 5B). The majority of the regulators that we found to be significantly activated or inhibited have been implicated into the control of HSC fate^34,35,36,37,38^. The most prominently inhibited regulators in *Apoe*^−/−^*Il27ra*^−/−^ LT-HSCs included Myc, CCND1 and E2F involved into proliferation; IFNα/β/ Ifnar, STAT1 and IL-27 directly or indirectly involved into IL-27R signaling pathways and control of HSCs expansion. On the other hand, proliferation-restrictive regulators including p53, mir-21, mir-223 and let-7^39,40,41,42,43,44^ were strongly activated in the absence of IL-27R signaling in LT-HSCs (Figure 5B). Genes whose expression is controlled by aforementioned regulators in LT-HSCs of *Apoe*^−/−^*Il27ra*^−/−^ mice are shown in a heatmap (p<0.05) (Figure 5C). Notably, among “activated regulators”, p53 had the most number of overrepresented targets among the gene list in LT-HSCs of *Apoe*^−/−^*Il27ra*^−/−^ mice (29 targets, p=2×10^−6^).

**Figure 5.**
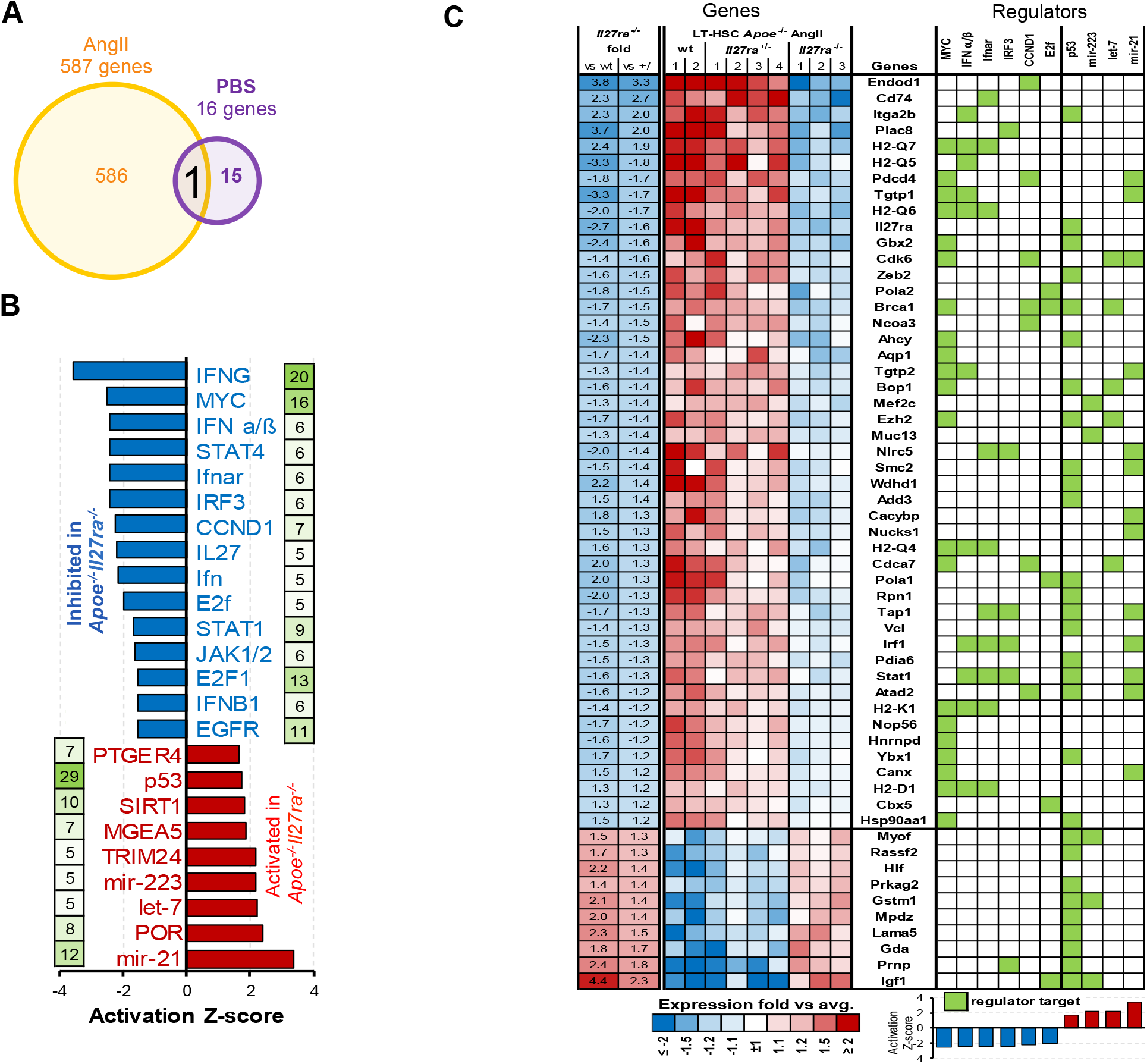
IL-27R signaling regulates the quiescence of HSCs in Ang II induced myelopoiesis. LT-HSCs (Sca-1^+^c-kit^+^CD150^+^CD48^−^) were FACS-sorted from bone marrow of *Apoe*^−/−^ (n=2), *Apoe*^−/−^*Il27ra*^+/−^ (n=4) or *Apoe*^−/−^*Il27ra*^−/−^ (n=3) mice fed with WD for 10wks and infused with Ang II or PBS for last 2 wks of feeding, followed by whole transriptome analysis. **(A)** Overall number of genes under Ang II and control PBS treatment affected by IL-27R deficiency. **(B)** De-regulated upstream regulators whose targets were significantly overrepresented among genes uniquely affected in IL-27R deficient mice. **(C)** Expression heatmap for significantly changed genes participating in regulators pathway. Green squares indicate known regulator->target relationship from published literature as recorded in Ingenuity Knowledgebase. *p<0.05.

These results suggest that IL-27R signaling maintains the balance between quiescence, proliferation and differentiation in LT-HSCs and show that the responsiveness of HSPCs to Ang II is blunted in the absence of IL-27R, suggesting that the lack of IL-27R signaling is associated with a previously described more quiescent state^45^. These data are consistent with our finding showing reduced BrDU incorporation in IL-27R deficient LT-HSCs. We next validated some of the genes from RNA sequencing analysis on isolated Lin^−^ HSPCs. Q-RT-PCR analysis revealed in IL-27R deficient cells a reduction of the expression of genes previously involved into regulation of proliferation and myeloid lineages differentiation^34,38,46,47^ (Supplemental Figure S6)

Along with the quiescence gene signature, IL-27R deficient HSPCs from Ang II treated mice were characterized by elevated p21 (Figure 6A). Importantly, infusion of Ang II into *Apoe*^−/−^ *Il27ra*^+/−^ controls triggered p21 downregulation, whereas p21 was still expressed in *Apoe*^−/−^*Il27ra*^−/−^ HSPCs (Figure 6A). To assess whether the observed effect on p21 expression was dependent upon Ang II and not another aspect of the BM microenvironment, we isolated HSPCs from *Apoe*^−/−^*Il27ra*^+/−^ or *Apoe*^−/−^*ll27ra*^−/−^ mice and stimulated them *in vitro* with Ang II. We found that direct Ang II stimulation strongly reduces p21 expression in control *Apoe*^−/−^*Il27ra*^+/−^ cells, but not in *Apoe*^−/−^*Il27ra*^−/−^ HSPCs (Figure 6B).

**Figure 6.**
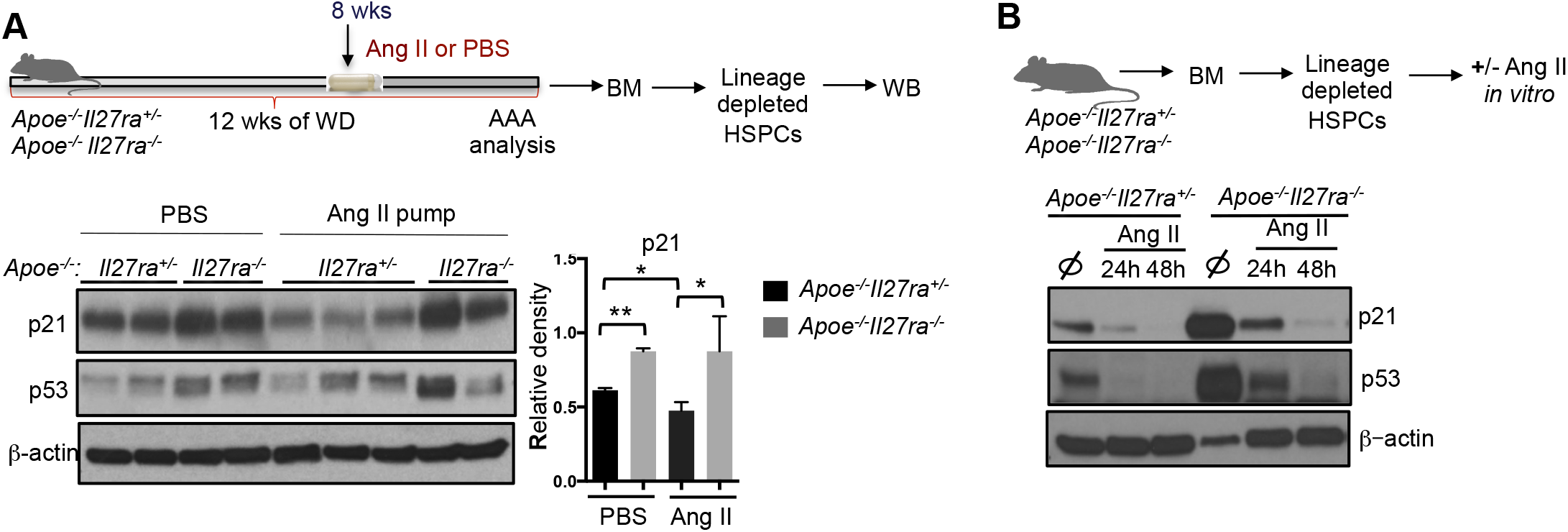
Increased expression of p21 in HSPCs of IL-27R deficient mice. **(A)** Lin^−^ HSPCs isolated from WD-fed *Apoe*^−/−^*Il27ra*^+/−^ or *Apoe*^−/−^*Il27ra*^−/−^ mice infused with PBS or Ang II for 4 weeks were lysed and subjected to western blotting with p21, p53 and b-actin antibody. *p<0.05, **p<0.01 **(B)** WB on Lin^−^ HSPCs sorted from naive *Apoe*^−/−^*Il27ra*^+/−^ or *Apoe*^−/−^*Il27ra*^−/−^ mice and stimulated *in vitro* with AngII for 24 or 48 hours.

Several signaling pathways could participate in the regulation of p21 expression, including interferon and Myc pathways, both of which are enriched in Ang II-treated wild type cells (Figure 5B, C). In addition, p53 is one of the key upstream bona fide regulators of p21 expression. p53 displayed the most number of overrepresented targets among the gene list (29 targets, p=2×10^−6^; z-score=1.74). We also found that p53 protein expression was reduced by *in vitro* stimulation with Ang II in *Apoe*^−/−^*Il27ra*^+/−^ HSPCs, while the downregulation was incomplete in IL-27R deficient cells (Figure 6B).

Collectively, these data suggest that HSPCs depends on IL-27R signaling to regulate the expression of p21 and other quiescence/proliferation regulators. Therefore, IL-27R signaling is required to control the balance between quiescence and cell-cycle entry in response to AngII - induced myelopoiesis during AAA progression (Figure 7).

**Figure 7.**
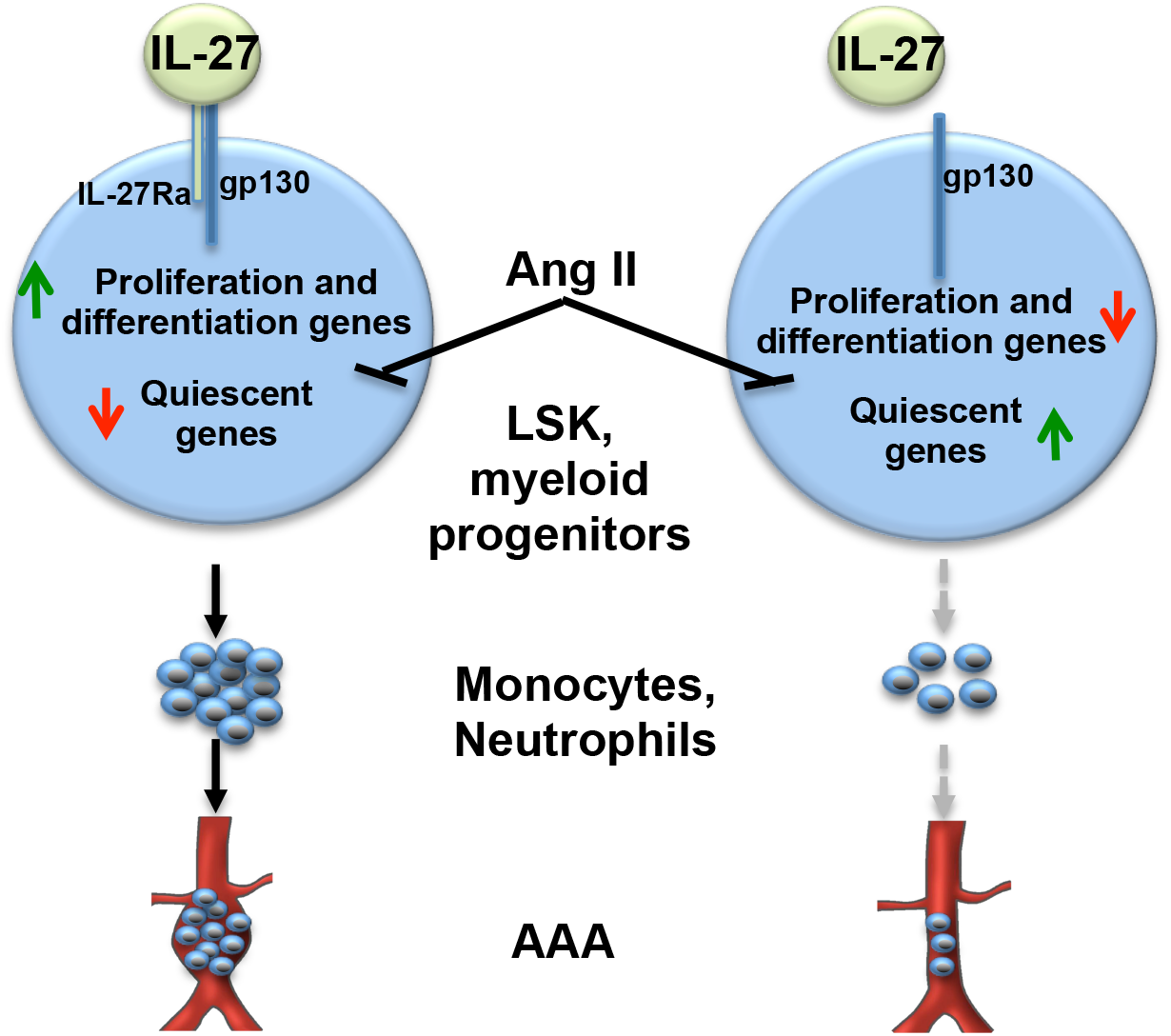
Role of IL-27R signaling in regulation of Ang II-induced myelopoiesis and AAA. Elevation of Ang II in AAA provides a stimulus for the activation of “stress” myelopoiesis via regulation of gene expression controlling proliferation and differentiation in IL-27R sufficient LT-HSCs. Lack of IL-27R signaling renders AngII-primed cells unable to overcome quiescent status, significantly reduces proliferation of progenitors and myeloid cell bone marrow output, reduces cell recruitment to AAA lesions and blocks AAA progression.

## Discussion

The function of IL-27R signaling was extensively investigated in various infectious models^23,24,25^, and an anti-inflammatory role IL-27R has been demonstrated in atherosclerosis^26,27,28^. Moreover, some of the IL-27R deficient mice employed in the atherosclerosis studies developed frank abdominal aorta lesions, similar to incipient AAA sites, raising the possibility that IL-27R might also restrict spontaneous AAA. The role of IL-27R in AAA, another vascular pathology with clear role of yet unidentified immune cytokine-driven mechanisms, however has never been assessed. Here we made the unanticipated observation that *Apoe*^−/−^*Il27ra*^−/−^ mice were largely protected from Ang II-induced AAA. This correlated with diminished accumulation of monocytes and neutrophils in suprarenal aortas, which is a key early event in AAA^13,14,15,16,17,41^. The expression of myeloid cell derived cytokines and chemokines was also decreased in the AAA lesions of *Apoe*^−/−^*Il27ra*^−/−^ mice infused with Ang II. Although homing mechanisms play an important role in the regulation of immune cell accumulation at the site of inflammation (AAA)^17,48^, the recruitment of a large amount of immune cells into rapidly developing AAA lesions should also require an increased BM output in production of these cells, known as “stress-induced” BM myelopoiesis. The unifying regulators, required for the AAA development, vessel inflammation and “stress myelopoiesis”, however remained unknown. Ang II was shown to induce the mobilization of splenic monocytes, while splenectomy and subsequent reduction of circulating monocytes suppresses AAA development^16^. Moreover, Ang II was shown to act directly on HSCs in BM and induce their self-renewal and rapid amplification of myeloid progenitors with subsequent production of mature myeloid cells^22^. Here we found that in our model Ang II causes the expansion of various HSCs and progenitor cells, including LT-HSCs and CD48^+^CD150^−^ population as well as more mature myeloid precursors in the BM of control *Apoe*^−/−^ or *Apoe*^−/−^*Il27ra*^+/−^ mice. A surprising observation of our studies was, however, that inactivation of IL-27R signaling significantly blunted Ang II-driven expansion of HSCs and progenitor cells, indicating a novel role of IL-27R signaling potentiating Ang II-induced myelopoiesis, which in turn promotes AAA.

Several cytokines have been implicated in the regulation of immune cells in AAA site^3,29,49,50,51,52,53,54,55^; however, the possible contribution of cytokine signaling in controlling of AAA-driven “stress induced” HSCs expansion and differentiation toward myeloid lineages has never been reported. Here, we for the first time establish IL-27R signaling as a critical regulator of Ang II-driven proliferation and differentiation of HSC, essential for Ang II-induced stress myelopoiesis in a non-infectious, chronic vascular injury model. Given the role of myeloid cells in host defense and tissue repair^56,57,58^ this cytokine-driven induction of HSCs may be a common mechanism regulating the rapid need for increased BM output during various pathophysiological processes.

Quiescence is one of the key characteristics of HSCs at the steady state, when a significant BM output of myeloid cells is not needed. HSCs quiescence is tightly regulated by many factors including IFN signaling and miRNAs (mir-21, mir-223, let-7)^36,59,60,61^. Maintenance of quiescence is crucial for prevention of both stem cell pool exhaustion and development of hematopoietic malignancies^62,63^. In chronic diseases, like AAA, prolonged exposure to “stress” factors ultimately affecting BM cells, such as high fat diet and Ang II^22,64^, may provoke HSCs to awaken from quiescence and activate stress myelopoiesis. Here we establish IL-27R signaling as an *in vivo* regulator of HSCs function in non-infection model of vascular injury and AAA. This role of IL-27 is in line with the studies where forced transgenic overexpression of IL-27 causes myeloproliferation in mice and malaria infection drives bone marrow alterations in IFN/IL-27 dependent manner^65,66^.

By performing RNAseq analysis of purified LT-HSCs we found that IL-27R deficiency rendered LT-HSCs unable to upregulate the expression of genes involved in cell proliferation and myeloid cell differentiation normally induced by Ang II. Moreover, IL-27R deficient LT-HSCs were characterized by a transcriptional profile that supports a quiescent state even following exposure to Ang II. Specifically, we found that ablation of IL-27R causes the upregulation of mir-21, mir-223, let-7 and p53 and the downregulation of E2F, CCND1, IRF3, IFN-“gene signature” and Myc, which are have been linked to a less proliferative and more quiescent state. In agreement, we also detected reduced BrDU incorporation in *Apoe*^−/−^*Il27ra*^−/−^ HSCs *in vivo*. Many of these regulators have also been previously suggested into the control HSCs fate. Indeed, mir-21 was previously implicated in hematopoietic suppression via activation of TGFβ signaling^39^. Mir-223 has been reported to attenuate hematopoietic cell proliferation^40^, while let-7 was shown to regulate HSCs fate by controlling self-renewal, proliferation, quiescence and differentiation via inhibition of TGFβ pathway^67,68^. Moreover, let-7 family members were shown to repress cell cycle regulators (Cyclin D1) and negatively regulate Myc expression^42,43,44^. Our RNA seq data also revealed downregulation of Cyclin D1, E2F, Myc and “IFN-gene signature”. TGFβ, p53 and IFN pathways may converge at the level of p21 expression, a key negative regulator of cell proliferation, whose levels are decreased by Ang II stimulation in an IL-27R dependent manner. Taken together, the gene expression profile of IL-27R-deficient LT-HSCs is characteristic of quiescent state and remains as such even when AAA-inducing AngII driven stress is applied.

Collectively, our data establish an unexpected role of IL-27R signaling in AAA and suggest that Ang II, a stress factor that is constantly present in AAA, perhaps, in combination with other factors like HFD, promotes HSPCs proliferation and reveals a previously unexplored requirement for IL-27R signaling in Ang II-driven HSCs expansion and myelopoiesis during AAA development. Our data also identifies IL-27 as a potential testable target for prevention and treatment of AAA and other forms of vascular injury, which require HSCs BM mobilization for its full pathology. Therefore, overstimulation of this otherwise anti-inflammatory signaling pathway can actually be pathogenic in AAA due to its ability modulate hematopoiesis.

## Methods

### Mice

*Il27ra*^−/−^ (*JAX* #018078) mice were crossed to *Apoe*^−/−^(*JAX* # 002052) mice to obtain *Apoe*^−/−^, *Apoe*^−/−^*Il27ra*^+/−^ and *Apoe*^−/−^*Il27ra*^−/−^ mice. C57BL/6 CD45.1 (JAX#002014) mice were crossed to *Apoe*^−/−^ mice to obtain CD45.1 *Apoe*^−/−^ mice. All mice were on C57BL/6 background. Mice were bred and housed under specific pathogen-free conditions in an AAALAC-approved barrier facility at FCCC. The genotyping was performed by standard PCR protocols. Animal numbers for each specific analysis are given in the Figure legends. *Apoe*^−/−^, *Apoe*^−/−^*Il27ra*^+/−^ and *Apoe*^−/−^ *Il27ra*^−/−^ mice were fed with “Western Diet” (WD) (Teklad 88137) for 8 weeks beginning at 8 weeks after birth followed by Ang II containing pump subcutaneous implantation. 4 weeks later mice were sacrificed and abdominal aortic aneurysm formation was analyzed. All animal experiments were approved by the Animal Care Committee at FCCC.

### implantation of Angiotensin II pumps

Ang II containing osmotic mini-pumps were prepared and implanted as previously described^5^. Briefly, mice were anesthetized and osmotic mini-pumps (Alzet 2004) loaded with Angiotensin II (800ng/kg/min; Calbiochem) were surgically subcutaneously implanted in the mid-scapular area over the shoulder blade followed by closing the wound with clips. Abdominal aortic aneurysm formation was analyzed after 14 or 28 days of Angiotensin II infusion.

### Biood pressure measurements

Systolic blood pressure was measured on conscious mice after 4 weeks after infusion of Ang II using tail cuff systems (CODA Non-Invasive Blood Pressure Monitor, Kent Scientific Corporation, Torrington, CT) according to manufacture’s protocol. Briefly, mice were placed in the holder maintained on the Animal Warming Platform. “Occlusion Cuff” and “VPR Cuff” were placed near the base of the tail. The cuffs were attached to the CODA Controller. 20 cycles of measurement were performed and accepted cycles were automatically displayed in spreadsheet format within the CODA application. Data were process in Excel by calculation of the average and standard deviation.

### Histology and immunofluorescence

For histological analysis, tissue of suprarenal aortas with or without abdominal aortic aneurysm were isolated and embedded in Tissue-Tek O.C.T. (Optimal Cutting Temperature) compound (Akura Finetechnical Co. Ltd., Tokyo, Japan) and stored at −80°C. 5-μm serial sections were cut and for representative images the sections were stained with hematoxylin (RICCA, Arlington, TX) and eosin (ThermoScientific, Kalamazoo, MI) (H&E staining). All images were acquired with microscope Eclipse 80i.

Immunofluorescence staining was performed as previously describe^27^. Briefly, 5-μm frozen sections of suprarenal aortas, containing abdominal aortic aneurysm, were fixed in cold acetone for 10 min at room temperature followed by fixation in 1% paraformaldehyde in 100mmol/L dibasic sodium phosphate containing 60 mmol/L lysine and 7 mmol/L sodium periodate at pH 7.4 on ice. After sections were blocked with avidin/biotin (Vector Laboratories, Burlingame, CA) for 10 min each, followed by blocking with 5% normal goat serum in 1% BSA in PBS for 15 min. Sections were stained at 4°C overnight with primary rat anti-mouse CD11b-FITC (M1/80; BD Bioscience) and rat anti-mouse Ly6G-FITC (1A8; Biolegend) antibody, followed by staining with secondary antibody for 1 hour at room temperature: goat anti-FITC AlexaFluor 488 (Molecular Probes), goat anti-rat Alexa Fluor 568 (Molecular Probes). Sections were stained with DAPI and mounted with Prolong Gold (Life Technologies, Eugene, OR). Images were examined on Leica SP8 DM6000 confocal microscope using HCX PLADO 20x and 40x oil-immersion objectives at 488, and 563 nm excitation wavelengths.

### Verhoeff-Van Gieson staining of elastic fibers

5-μm sections of suprarenal aortas (with or without AAA) were cut and staining of elastic fibers was performed. Frozen sections were hydrated followed by staining in Verhoeff’s solution for 1h. After slides were differentiated in 2% ferric chloride for 2 min and treated with 5% sodium thiosulfate for 1 min. Sections were counterstained in Van Gieson’s solution for 5 min and dehydrated (all reagents were purchased from Electron Microscopy Sciences, Hatfield, PA). All images were acquired with microscope Eclipse 80i.

### Flow cytometry analysis of isolated cells from AAA

Cells were isolated from suprarenal aortas (with or without AAA) of *Apoe*^−/−^*Il27ra*^+/−^ and *Apoe*^−/−^ *Il27ra*^−/−^ mice infused with Ang II and analyzed by flow cytometry. Briefly, mice were sacrificed by CO_2_ inhalation, and aortas were perfused with PBS containing 2% heparin to remove all traces of blood. Suprarenal aortas were isolated, cut into small pieces and incubated in a cocktail of digestion enzymes containing hyaluronidase (120 U/ml), collagenase I (450 U/ml), collagenase XI (250 U/ml) and DNAse I (120 U/ml) (all enzymes were from Sigma, St Louis, MO) for 55 min at 37°C with gentle shaking. After incubation cell suspensions were filtered through 70-μm cell strainer and stained with CD45-PerCP (30-F11; Biolegend), CD11b-PacBlue (M1/70; eBioscience), CD11c-Cy7PE (N418; eBioscience), Ly6G-AF780 (1A8; Biolegend), Ly6C-FITC (HK1.4; Biolegend) and LIVE/DEAD Fixable Yellow Dead Cell dye (Lifetechnologies, Eugene, OR) for flow cytometry (BD LSR II). Data were analyzed using Flowjo Software (Version 9.7.6).

### Flow cytometry analysis of HSPCs

Bones (femur/tibia) were isolated from *Apoe*^−/−^*Il27ra*^+/−^ and *Apoe*^−/−^*Il27ra*^−/−^ mice implanted with Ang II or PBS-containing pumps for 4 weeks. Single cell suspension was filtered via 70-μm cell strainer and RBC were lysed by RBC Lysis Buffer. Cells were stained for HSPCs subpopulation markers and analyzed by Flow cytometry (BD LSR II). Live cells were defined as LIVE/DEAD negative. CD3-Cy5PE (145-2c11; Biolegend), CD4-Cy5PE (RM4-5; Biolegend), CD8a-Cy5PE (53-6.7; Biolegend), CD19-Cy5PE (6D5; Biolegend), B220-Cy5PE (RA3-6B2; Biolegend), Gr1-Cy5PE (RB6-8C5; Biolegend) and Ter119-Cy5PE (TER-119; Biolegend) antibody were used to define mature lineages. HSPCs were stained using c-Kit-APC (2B8; BD Biosciences), Sca-1-Cy7PE (D7; eBioscience), CD150-PE (TC15-12F12.2; Biolegend), CD34-FITC (RAM34; BD Biosciences), FcgRII/III (CD16/32)-Cy7APC (93; eBioscience) and CD48-PacBlue (HM48-1; Biolegend). Cell sorting was performed on FACS Aria II cell sorter (BD Bioscience).

### in vivo BrDU incorporation

Mice were injected i.p. with a single dose of BrDU (1mg/mouse) (BD Pharmigen) in sterile PBS. Bones were harvested 4 hours after injection. BrdU incorporation was assessed by Flow cytometry (BD LSR II) using BrDU Flow Kit (BD Pharmingen) according to the manufacture’s protocols. Data were analyzed using Flowjo Software (Version 9.7.6). Briefly, BM cells were stained for HSPCs subpopulation markers, followed by fixation in BD Cytofix/Cytoperm Buffer for 20 min and in BD Cytoperm Plus Buffer for 10 min on ice. Next, cells were re-fixed in BD Cytofix/Cytoperm Buffer for 5 min on ice and treated with DNase to expose incorporated BrDU for 1 hour at 37°C. Cells were stained with anti-BrDU-PerCP Ab for 20 min at RT. LIVE/DEAD staining was used to exclude dead cells.

### immunomagnetic isolation of HSPCs

HSPCs were isolated using Easy Sep Mouse Hematopoietic Progenitor Cell Isolation Kit (StemCell Technologies) according to manufactures’s protocol. Briefly, BM cells from *Apoe*^−/−^ *Il27ra*^+/−^ and *Apoe*^−/−^*Il27ra*^−/−^ mice were incubated for 15 min at 4°C with hematopoietic progenitor cells biotin isolation cocktails followed by incubation with Streptavidin RapidSpheres for 10 min at 4°C. Tubes with cell suspension were placed into the magnet and incubated for 4 min followed by pouring the enriched cell suspensions. Collected fraction represented the enriched lineage negative cell fraction (HSPCs). After isolation, Lin^−^ HSPCs were counted and used for gene expression analyses or Colony Formation Assay.

### Western Blot

Cell lysates of sorted HSPCs were separated by 4-12% Bis-Tris Protein Gels (Invitrogen) and transferred to PVDF transfer membranes (Invitrogen). Each membrane was washed with TBST (10 mM Tris-HCl (pH 7.6), 150 mM NaCl, 0.05% Tween-20) and blocked with 5% skimmed milk for 1 h prior to incubation with a 1:1,000 dilution of the appropriate primary antibody. Each membrane was washed, and primary antibodies were detected with a 1:5000 dilution of HRP-conjugated rabbit anti-mouse IgG or mouse anti-rabbit IgG (Cell Signaling). The reactive bands were visualized with enhanced KODAK chemiluminescence BioMax film (Carestream Health Inc.).

### Colony Formation Assay

Isolated HSPCs from *Apoe*^−/−^*Il27ra*^+/−^ and *Apoe*^−/−^*Il27ra*^−/−^ mice infused with Ang II were sorted and plated at 1×10^3^ cells in 1ml of MethoCult^®^ GF M3434 (Stem Cell Technologies) according to manufacturer’s instructions. Serial re-plating was performed at day 6 and the cells were cultured until day 16. Total numbers of colonies were counted at day 6 and 16 under the light microscope (Nikon TMS).

### Competitive BM reconstitution assay

*Apoe*^−/−^ CD45.1 recipient mice were irradiated in 2 doses of 550 rad each (for a total 1100 rad), three hours apart. Femurs and tibias of donor mice (*Apoe*^−/−^ CD45.1 or *Apoe*^−/−^*Il27ra*^−/−^ CD45.2) were harvested, flashed in sterile conditions using PBS +2% FBS and filtered through 70-μm cell strainer. Donor mixes were prepared by mixing *Apoe*^−/−^ CD45.1 and *Apoe*^−/−^*Il27ra*^−/−^ CD45.2 BM cells in a 90%:10%, 10%:90% and 50%:50% ratio and resuspended in sterile PBS. 250 μL of total BM suspension, containing 5×10^6^ cells was i.v. injected into each irradiated recipient mouse. After BMT, recipient mice were provided with autoclaved acidified water, containing antibiotic (trimethoprim-sulfamethoxazole) for 2 weeks. 1 month after reconstitutuion, mice were placed on high fat diet “Western Diet” (WD) (Teklad 88137) for 8 weeks followed by AngII containing pump subcutaneous implantation. Mice were sacrificed 4 weeks after pump implantation. BM, spleen, blood, suprarenal aorta and arch were harvested and analyzed by FACS. The reconstitution efficiency was analyzed by FACS 4 weeks after BMT using CD45.1-APC (A20; eBioscience) and CD45.2-PerCP (104; Biolegend) antibody.

### RNA-sequencing and data processing and analysis

A total of 5,000 LT-HSCs (Sca-1^+^c-kit^+^CD150^+^CD48^−^) were FACS-sorted from the BM of *Apoe*^−/^, *Apoe*^−/−^*Il27ra*^+/−^ or *Apoe*^−/−^*Il27ra*^−/−^ mice infused with Ang II or PBS for 2 weeks. RNA was extracted using Quigen RNA isolation kit according to manufacture protocol. Total RNA libraries were prepared by using Pico Input SMARTer Stranded Total RNA-Seq Kit (Takara). 250pg-10ng total RNA from each sample was reverse-transcribed via random priming and reverse transcriptase. Full-length cDNA was obtained with SMART (Switching Mechanism At 5’ end of RNA Template) technology. The template-switching reaction keeps the strand orientation of the RNA. The ribosomal cDNA is hybridized to mammalian-specific R-Probes and then cleaved by ZapR. Libraries containing Illumina adapter with TruSeq HT indexes were subsequently pooled and loaded to the Hiseq 2500. Paired end reads at 75bp with 30 million reads per sample were generated for the bioinformatic analysis. RNA-seq data was aligned using STAR^69^ against mm10 genome and RSEM v1.2.12 software^70^ was used to estimate gene-level read counts using Ensemble transcriptome information. Only samples with at least 20% exonic reads considered by RSEM were used for analysis. DESeq2^71^ was used to estimate significance of differential expression difference between any two experimental groups with mouse gender used as an additional factor. Overall gene expression changes were considered significant if passed FDR<5% thresholds. Gene set enrichment analysis was done using QIAGEN’s Ingenuity^®^ Pathway Analysis software (IPA^®^, QIAGEN Redwood City,www.qiagen.com/ingenuity) on genes that passed nominal p<0.05 using “Canonical Pathways” and “Upstream Regulators” options. Only pathways that passed FDR<10% threshold and had all genes de-regulated in the same direction were considered. Upstream regulators with significantly predicted activation state (|Z-score|>1.5) that in addition passed p<0.05 target enrichment threshold with at least 5 target genes were reported.

### RNA isolation and Gene Expression

Suprarenal aortas /AAA were isolated from *Apoe*^−/−^*Il27ra*^+/−^ and *Apoe*^−/−^*Il27ra*^−/−^ mice implanted with Angiotensin II pumps and homogenized in PureZol (Bio-Rad) with Ceramic Beads using Omni Bead Ruptor 24. Total RNA was extracted using Aurum^™^ Total RNA Fatty and Fibrous Tissue Kit (Bio-Rad) according to the manufactures’s instructions. HSPCs were lysed in RLT Plus buffer and total RNA were isolated using RNeasy Plus Mini Kit (Qiagen) according to the manufactures’s protocols. cDNA was synthesized using iScript Reverse Transcription Supermix™ (Bio-Rad) with random primers according to the manufacters’s protocol. qRT-PCR was performed with the Cycler CFX 96 connect Real Time System (Biorad) using iTaq Universal SYBR Green Supermix (BioRad). The following primers were used: ribosomal L32, CCL2, CCL5, CCL22, MMP-9, MMP-2, TNF-α, IL-1β, MPO, IL-3Ra, CSFR3R, KLF5, c-Myc, Cyclin D. Sequences of the primers were obtained from National Institutes of Health qPrimerDepot (http://pga.mgh.harvard.edu/primerbank/). Gene expression was normalized to ribosomal L32 expression.

### Statistics

Student’s 2-tailed *t* test was used to compare conditions. Data analyses were performed using the GraphPad Prism Software. Data are presented as mean ± SEM; *p<0.05, **p<0.01, ***p<0.001. p<0.05 was considered statistically significant.

## Acknowledgments

We acknowledge the help of FCCC Facilities as well as Bioinformatics facility at The Wistar Institute. We thank Dr. Sergei Grivennikov (FCCC) and Dr. David L. Wiest (FCCC) for critical reading of the manuscript.

This work was supported NCI P30 Cancer Center Grant to FCCC; and by AHA SDG 13SDG14490059, WW Smith Charitable Trust, NIH R21 CA202396, NIH R56 HL133669 and NIH R01 HL133669 grants to E.K.K.

## Authorship Contributions

IIP, TA and EKK designed the study and planned the experiments; IIP, TA and ARF performed the experiments. IIP, TA and EKK wrote the manuscript. SMS provided assistance with data interpretation and manuscript writing. SE provided help with critical reading of the manuscript. YFT and AK assisted with RNA-seq analysis.

