## Supplementary figures for "IL-27 receptor signaling regulated stress myelopoiesis drives Abdominal Aortic Aneurysm development"

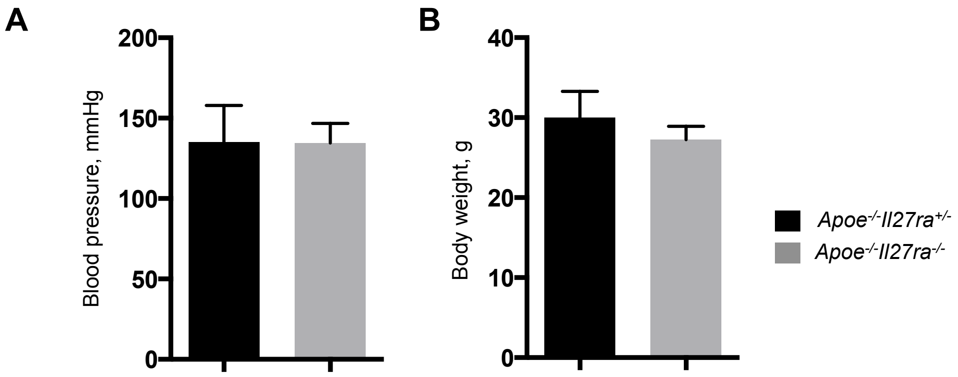


**Supplemental Figure 1. IL-27R deficiency does not affect blood pressure and body weight. (A**) Systolic blood pressure was measured on conscious *Apoe*$^{-/-}$*Il27ra*$^{+/-}$ (n=8) and *Apoe*$^{-/-}$*Il27ra*$^{-/-}$ (n=5) mice infused with Ang **II** for 4 weeks using tail cuff system. (**B**) Body weight of *Apoe*$^{-/-}$*Il27ra*$^{+/-}$ (n=6) or *Apoe*$^{-/-}$*Il27ra*$^{-/-}$ (n=6) mice after Ang **II** infusion.

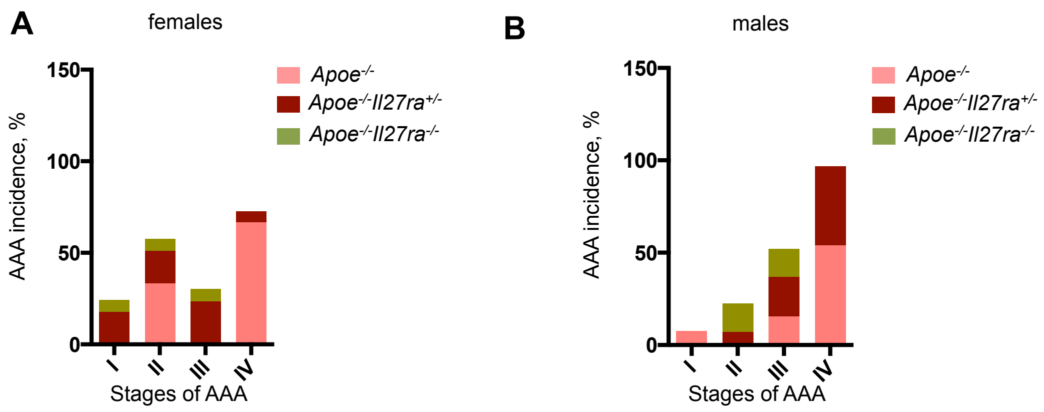


**Supplemental Figure 2.** Classification of AAA stages based on severity grade in female **(A)** and male **(B)** mice.

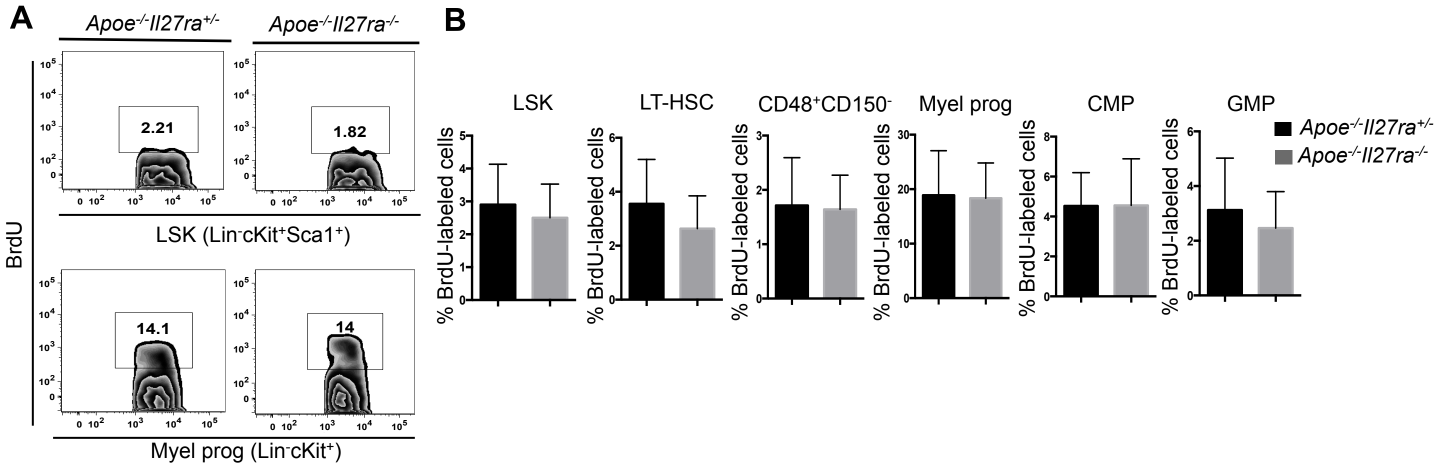


**Supplemental Figure 3. IL-27R deficiency does not affect proliferation of HSPCs in a steady state.** Proliferation of bone marrow HSPCs isolated from *Apoe$^{-/-}$Il27ra$^{+/-}$* (n=8) or *Apoe$^{-/-}$Il27ra$^{-/-}$* (n=6) mice fed with WD and infused with PBS as determined by BrdU incorporation. Representative dot plots (**A)** and percentage **(B)** of live BrdU positive LSK (Lin$^-$c-kit$^+$Sca1$^+$) and myeloid progenitors (Lin$^-$c-kit$^+$Sca1$^-$), including LT-HSC (CD150$^+$CD48$^-$), CD48$^+$CD150$^-$, CMP (Lin$^-$c-kit$^+$CD34$^+$FcgR$^-$) and GMP (Lin$^-$c-kit$^+$CD34$^+$FcgR$^+$). Data are mean ± SEM from 2 independent experiments.

**A**

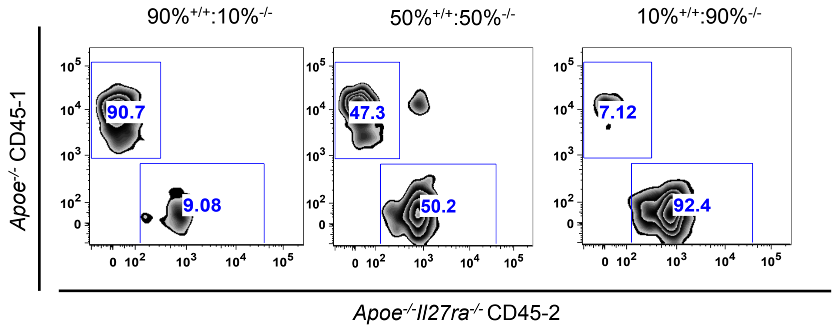


**Supplemental Figure 4.** Bone marrow reconstitution efficiency was determined by flow cytometry in the peripheral blood of recipient mice 4 weeks after competitive bone marrow transplantation.

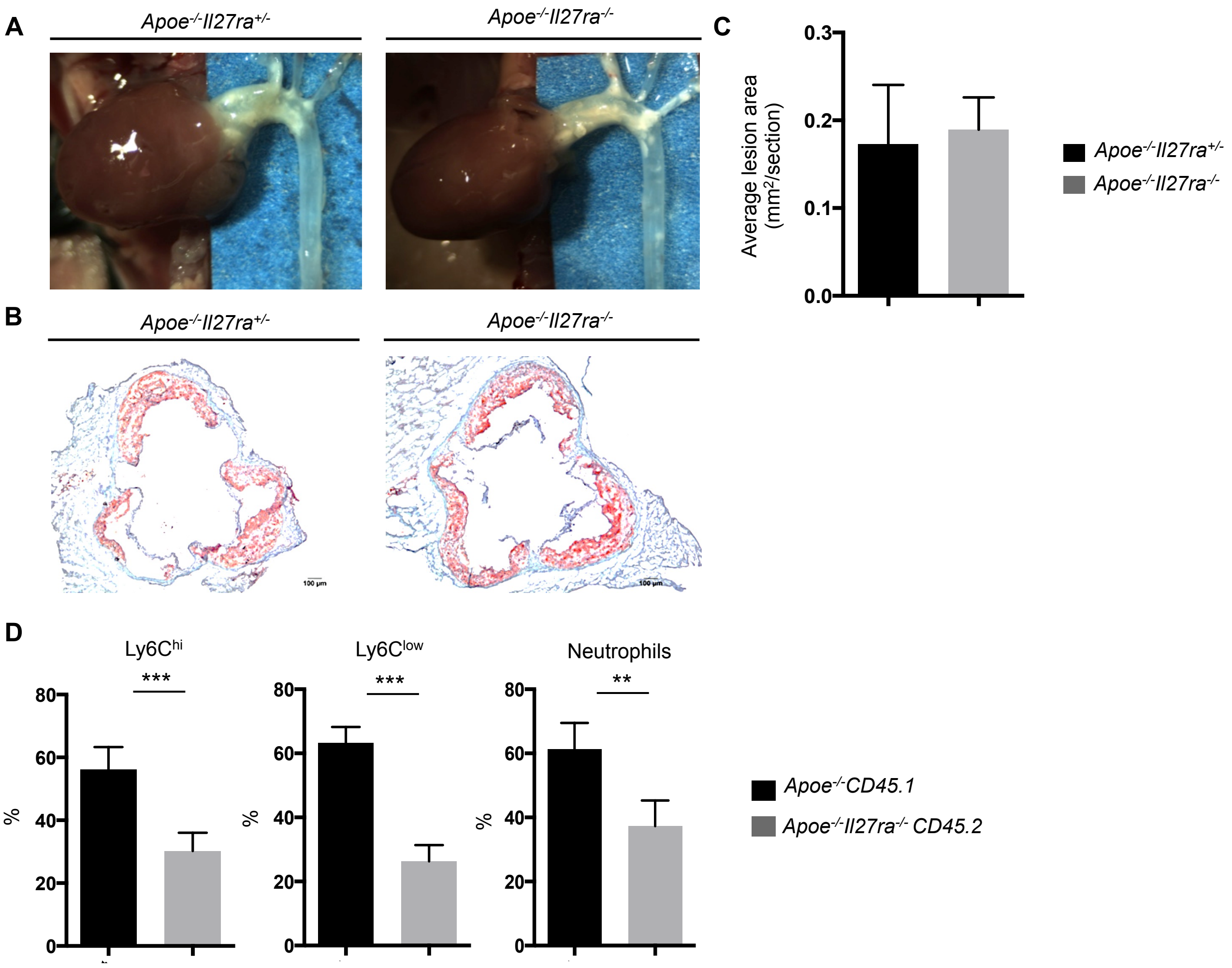


**Supplemental Figure 5. Ang II infusion strongly accelerates atherosclerosis development in IL-27R sufficient, but not deficient mice. (A)** Images of atherosclerotic lesions in aortic arch and **(B)** aortic root sections of *$Apoe^{-/-}$ $Il27ra^{+/-}$* and *$Apoe^{-/-}$ $Il27ra^{-/-}$* mice fed with WD for 12 weeks and infused with Ang II for last 4 weeks of feeding. **(C)** Quantitative comparison of aortic lesion size in *$Apoe^{-/-}Il27ra^{+/-}$* (n=5) and *$Apoe^{-/-}$ $Il27ra^{-/-}$* (n=5) mice. Data are mean ± SEM from 2 independent experiments. (**D**) Proportion of live donor-specific $Ly6C^{hi+}$, $Ly6C^{low+}$ monocytes and $Ly6G^{+}$ neutrophils in aortic arch of mice reconstituted with donor mixes of 50% *$Apoe^{-/-}$* CD45.1 and 50% *$Apoe^{-/-}Il27ra^{-/-}$* CD45.2 total bone marrow cells and fed with WD for 10 weeks, where last 2 weeks they were infused with Ang II. Data are mean ± SEM from 2 independent experiments. *p<0.05, **p<0.01, ***p<0.005.

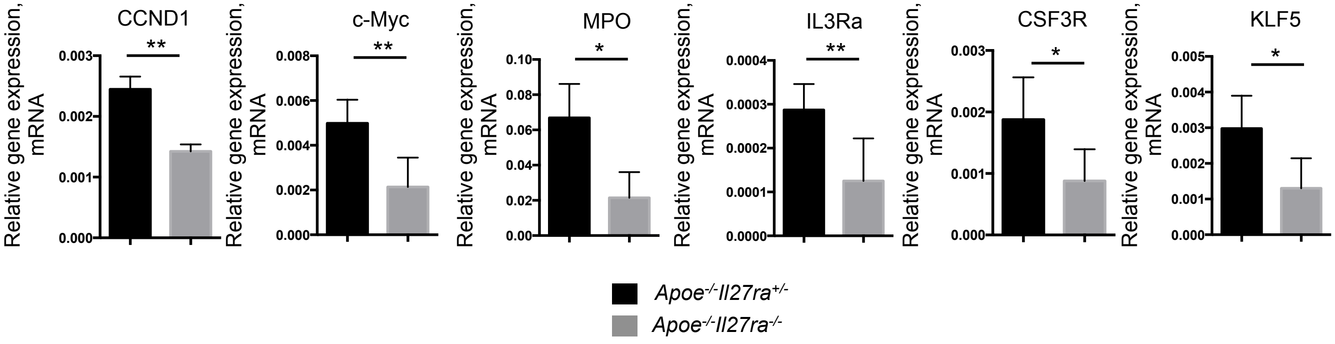


**Supplemental Figure 6. Decreased expression of genes regulating proliferation and myeloid lineage commitment in HSPCs of *Apoe$^{-/-}$Il27ra$^{-/-}$* mice.** Relative gene expression in Lin$^{-}$ HSPCs isolated from WD-fed *Apoe$^{-/-}$Il27ra$^{+/-}$* (n=5) or *Apoe$^{-/-}$Il27ra$^{-/-}$* (n=5) mice infused with Ang II for 4 weeks were normalized to L-32 gene expression **.** *p<0.05. **p<0.01. Data are mean ± SEM from at least 2 independent experiments.

| Canonical Pathway | *p-value* | FDR | Molecules specifically downregulated by Il27ra$^{-/-}$ |
|---|---|---|---|
| Antigen Presentation Pathway | $6x10^{-9}$ | 0.0% | NLRC5, CALR, HLA-A, CANX, CD74, TAP1, HLA-DRB5 |
| Th1 Pathway | 0.0003 | 2.5% | RUNX3, HLA-A, IL27RA, STAT1, HLA-DRB5, IRF1 |
| Unfolded protein response | 0.0005 | 2.5% | CALR, HSP90B1, PDIA6, CANX |
| Phagosome Maturation | 0.0005 | 2.5% | CALR, TUBA8, HLA-A, CANX, TAP1, HLA-DRB5 |
| Th1 and Th2 Activation Pathway | 0.0017 | 5.1% | RUNX3, HLA-A, IL27RA, STAT1, HLA-DRB5, IRF1 |
| Interferon Signaling | 0.0019 | 5.1% | STAT1, TAP1, IRF1 |
| Protein Ubiquitination Pathway | 0.0023 | 5.4% | HSP90B1, HLA-A, HSP90AA1, DNAJB6, PSMD4, BRCA1, TAP1 |

**Supplemental Table 1. Pathways de-regulated in *Apoe$^{-/-}$Il27ra$^{-/-}$* mice after Ang II infusion.** LT-HSCs (Sca-1$^{+}$ c-kit$^{+}$CD150$^{+}$CD48$^{-}$) were FACS-sorted from bone marrow of WD-fed *Apoe$^{-/-}$* (n=2), *Apoe$^{-/-}$Il27ra$^{+/-}$* (n=4) or *Apoe$^{-/-}$Il27ra$^{-/-}$* (n=3) mice infused with Ang II or PBS for 2 weeks, followed by whole transriptome analysis. Only pathways that passed FDR<10% threshold with the same de-regulation effect on all genes are reported.
